# Why mediation analysis trumps Mendelian randomization in population epigenomics studies of the Dutch Famine

**DOI:** 10.1101/362392

**Authors:** Elmar W. Tobi, Erik W. van Zwet, L.H. Lumey, Bastiaan T. Heijmans

**Affiliations:** Molecular Epidemiology, Department of Biomedical Data Sciences, Leiden University Medical Center, 2300 RC Leiden, The Netherlands; Medical Statistics, Department of Biomedical Data Sciences, Leiden University Medical Center, 2300 RC Leiden, The Netherlands; Department of Epidemiology, Mailman School of Public Health, Columbia University, New York, NY 10032, USA

## Abstract

Our recent analysis of genome-wide DNA methylation data in men and women exposed to the Dutch Famine met passionate criticism by several researchers active on Twitter. It also prompted a more reasoned letter by Richmond and colleagues. At the core of the debate is the proper interpretation of findings from a mediation analysis. We used this method to identify specific DNA methylation changes that statistically provide a link between prenatal exposure to famine and adult metabolic traits. Our critics first argue that our results do not suggest mediation but reverse-causation, where famine-induced metabolic traits drive changes in DNA methylation. We rebut this scenario in a simulation study showing that our test of mediation was unlikely to become statistically significant in the case of reverse-causation. Some critics then argue that Mendelian randomization provides the sole path to correct inference. This belief misses a crucial point: DNA methylation, especially when measured in peripheral blood, is not likely to be a causal mediator from a biological point of view. It could however be a proxy of epigenetic regulation changes in specific tissues, for example at the level of transcription factor binding. If so, a Mendelian randomization approach using genetic variants affecting local DNA methylation in blood will be disconnected from the underlying biological mechanism and is bound to yield false-negative results. Our new simulation studies strengthen the original reasoning that the relationship between prenatal famine and metabolic traits is statistically mediated by specific DNA methylation changes while the specific molecular mechanism awaits elucidation.

## Introduction

Intrauterine exposure to an adverse environment has been linked to adult health^1-3^ and epigenetic mechanisms are widely believed to mediate this long-term effect.^4,5^ Analyses aimed at finding the specific epigenetic changes involved are a logical next step. Previous analyses have concentrated on the association between intrauterine exposures and either DNA methylation or adult health, but not on both. As a result, mediating epigenetic changes remain ill-defined. We recently analyzed genome-wide DNA methylation data in men and women with intrauterine exposure to the Dutch Famine.^6^ We used a genome-wide search for potential mediators followed by statistical mediation analysis to identify specific genomic regions at which differential epigenetic regulation could explain the observed association of prenatal famine exposure with adult metabolic traits. Our analysis identified the DNA methylation level in blood at one CpG as a candidate mediator of the association between famine exposure and adult body-mass index (BMI), and at eight CpGs as candidate mediators of the association with adult serum triglycerides levels (TG).

In a response to our work, Richmond et al^7^ argue that not mediation but reverse-causation is the most likely explanation for our results. In their view, the observed DNA methylation differences are secondary to changes in BMI and TG induced by prenatal famine and do not drive the observed adult metabolic traits.

## Analysis

To obtain a quantitative view of the assumptions underlying the reasoning of Richmond et al. we carried out a series of simulation studies. We generated one thousand data sets according to both the mediation and the reverse-causation scenario using the data patterns as observed in our original study. We then recorded the fraction of significant results obtained by Sobel tests for mediation (computationally fast and valid, assuming that Monte Carlo procedures for estimating p-values are not critical for normally distributed simulated data). The scenarios and results of our simulations are summarized in Figure 1.

**Figure 1.**
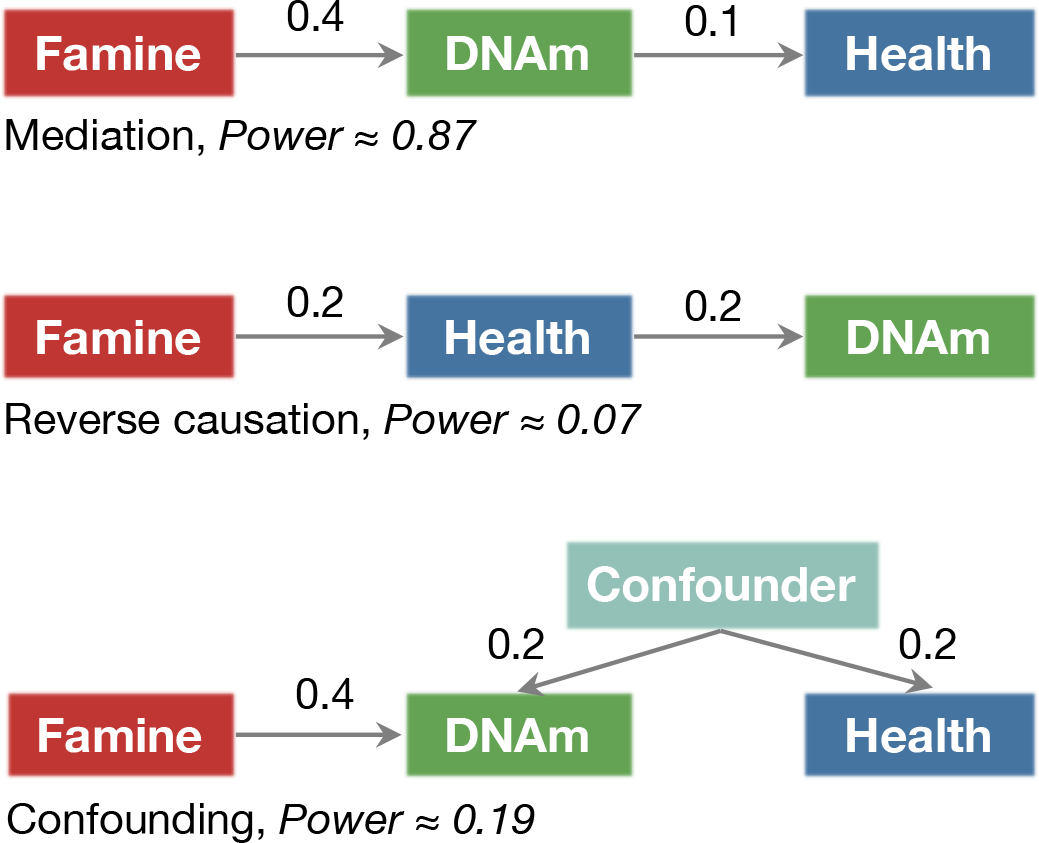
Statistical tests for mediation have low power for reverse-causation. A simulation study of three scenarios represented as directed acyclic graphs (DAGs) that could result in statistical evidence for mediation. We simulated 1000 data sets under the four scenarios and using realistic effect sizes estimated on the basis of empirical data from our study. We recorded the power of the Sobel test for mediation (α=0.05) for each scenario (depending on the scenario, statistical significance is interpreted as a true or false-positive finding). The test for mediation has high power if DNA methylation (DNAm) is on the path from prenatal famine (Famine) to metabolic traits (Health), ie. Mediation (Scenario A in Richmond et al^7^), but low power if changes in DNAm arise a secondary effect of famine-induced changes in health, ie. reverse-causation (Scenario B in Richmond et al), or if DNAm and Health are linked by an unobserved confounder confounding (Scenario D in Richmond et al).

Under the scenario of mediation, our simulations show a high statistical power of the mediation test (0.87). In contrast, the test for mediation has a very low power (0.07) under the reverse-causation scenario proposed by Richmond et al. This implies that the test for mediation is very unlikely to become statistically significant when changes in DNA methylation are secondary to changes in BMI and TG induced by prenatal famine. The findings from the simulation study agree with the results of our original analysis which effectively eliminated those CpGs as potential mediators that had been previously identified in epigenome-wide association studies of BMI and TG, but for which there was evidence of reverse-causation in 2-step Mendelian randomization analysis.^8,9^ Conversely, for the CpGs we identified as candidate mediators, there was no or inconclusive evidence for reverse-causation from previous studies.^8-10^ Richmond et al also considered a scenario in which DNA methylation and metabolic traits are linked by an unobserved confounder. The power for this scenario is low (0.19). In our original analysis was already adjusted for key potential confounders, including smoking, social-economic position, and current diet, further decreasing the likelihood of this scenario. Our new simulation study therefore effectively rebuts the comments of Richmond et al. and show that mediation by DNA-methylation changes is more likely to explain our findings than reverse-causation.

From a more general perspective however, we feel that neither of the above scenarios is likely to present a sufficiently accurate picture of the biological mechanism underlying our results. DNA methylation is known to mark the regulatory state of a genomic region but does not necessarily control that state. As an example, an extensive body of work has shown that DNA methylation changes can be the downstream effect of altered transcription factor binding.^11,12,13^ This suggests a scenario where the causal driver of the relationship between prenatal famine and metabolic traits could be a change in epigenetic regulation different from DNA methylation. Statistical evidence for mediation is observed if a CpG site is an adequate proxy of this causal change (Figure 2). This reasoning is not new and was already included in our mediation paper^6^ and also in earlier work.^5^ We have also simulated this driver scenario and found that a mediation analysis has considerable power to detect such effects (0.62).

**Figure 2.**
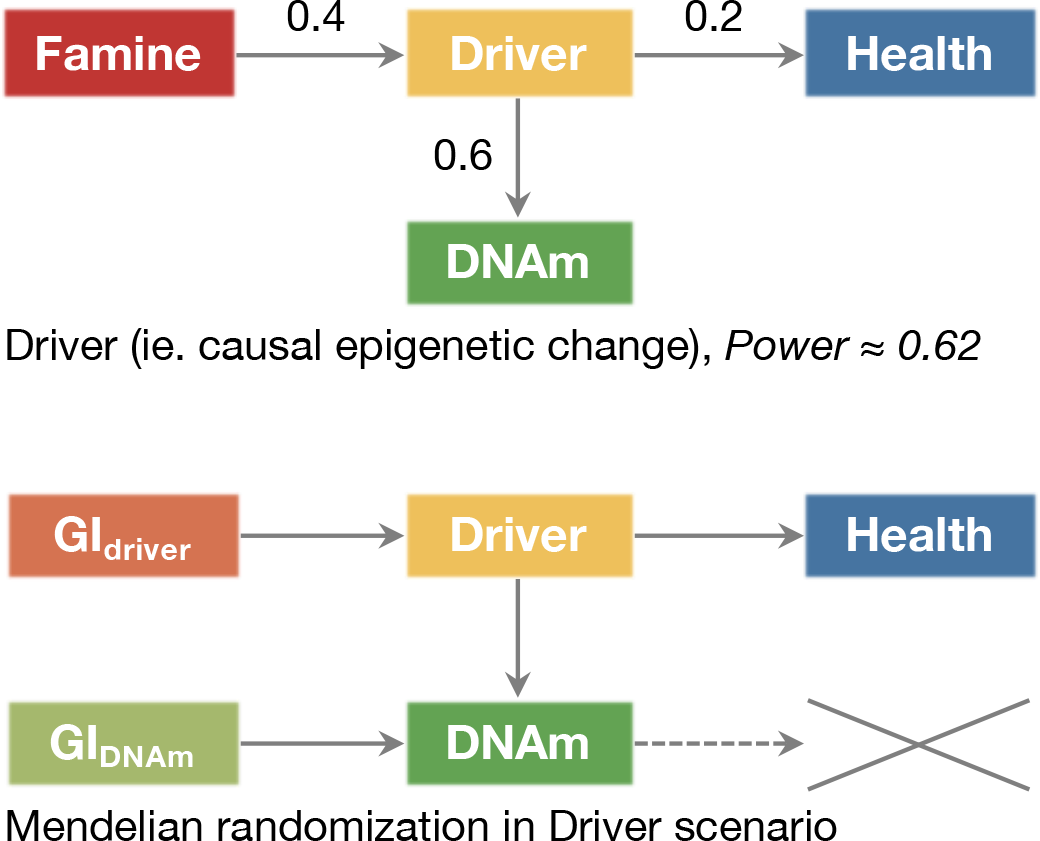
DNAm identified as mediator plausibly are proxies for the mechanistically causal change in epigenetic regulation. DNAm is known to frequently mark the epigenetic state of a region instead of control it. If DNAm is a proxy for another mechanistically causal change in epigenetic regulation that drives the relationship between Famine and Health, a test of mediation has power to detect such DNAm differences (akin to Scenario C in Richmond et al^7^). This scenario would reconcile the current limited success of Mendelian randomization approaches. Using a methylation QTL SNP as genetic instrument (GI) as done by Richmond et al will be turn out negative. A GI for the mechanistically causal change in epigenetic regulation is required to detect a causal effect on health.

The notion that DNA methylation in blood may not be the biological mediating mechanism has important implications for the application and interpretation of Mendelian randomization analysis. Richmond et al claim that Mendelian randomization is the only viable route to valid causal inferences in population epigenomics studies. This claim is problematic however, since current Mendelian randomization approaches use local genetic instruments to predict methylation of a CpG site in blood. The findings will therefore be negative in the event that DNA methylation change is merely a proxy of underlying causal changes in the epigenetic regulation at the level of histone modifications or transcription factor binding (Figure 2). This mismatch of unsuitable statistical methods to specific biological mechanisms may explain why there currently are surprisingly few examples of Mendelian randomization studies that link CpG methylation in blood to phenotypic changes.^10^ By contrast, there are ample examples of reverse-causation.^8-10^ It is therefore not surprising that Richmond et al failed to find evidence for a directed effect of methylation on BMI and TG for our candidate mediator CpGs. It is important to realize however that despite the disappointing results of Mendelian randomization approaches in this setting, few will dispute that epigenetic regulation could be an important driver of phenotypic variation.

## Conclusion

Our simulations confirm that mediation analysis is a legitimate and potentially valuable approach to identify potential causal mechanisms in population epigenomics studies. We also agree that Mendelian randomization can be a very powerful tool for causal inference in molecular data but any negative or positive findings will require cautious interpretation. Negative findings may be due to a mismatch between the statistical method and the underlying mechanism, and positive findings could be related to pleiotropy and backdoor paths.^8,14,15^ As in all modelling approaches to data analysis, any study result should be viewed in the broader context of the available knowledge from relevant disciplines and the relevance of study findings should not be determined by p-values from isolated statistical tests alone. In our original report^6^, we clearly outline the limitations of the Dutch Famine data and consider all available evidence to come to a cautious interpretation of our findings.

Identifying causal effects in observational data is challenging and results must be interpreted with caution.^16,17^ Causal reasoning in population epigenomics is best served by weighting evidence from alternative approaches. Essential in this field is the awareness that progress will depend on contributions from biology, epidemiology, and other disciplines. To identify molecular mechanisms in sufficient detail, population-based approaches (covering diverse study designs and analytical strategies) and experimental studies (from human cells and organoids to animal models) will be required from researchers who keep an open mind as to unexpected turns of biology and to optimal analytic methods to deal with the observed phenomena.

## Simulation script

To help further discussions on mediation analysis based on empirical data, we provide a set of R Markdown scripts to perform further simulations through GitHub (https://github.com/molepi/SimulationStudy-Mediation).

